# Circadian rhythmicity and photobiological mechanisms of light sensitivity and discomfort glare in humans

**DOI:** 10.1101/2024.02.05.578180

**Authors:** I Daguet, V Raverot, C Gronfier

## Abstract

Discomfort glare is a common visual sensation, which is generally reported when being exposed to a brighter lit environment. In certain clinical conditions, this sensation is abnormally amplified, and is commonly named photophobia. Despite the frequent appearance of this sensation in everyday life or in pathological conditions, the underlying mechanisms remain poorly understood. We show here, in highly controlled laboratory constant routine conditions, that light-induced discomfort glare is rhythmic over the 24-hour day. We reveal a strong circadian drive, with a sinusoidal rhythmicity, with maximal discomfort glare in the middle of the night and minimal in the afternoon. We also find a modest sleep-related homeostatic drive of visual discomfort, with a linear increase in discomfort glare over 34 hours of prolonged wakefulness. Our study reveals that discomfort glare is primarily driven by the ipRGC pathway, and that mid and/or long wavelengths cones are involved as well. The 6.5-hour phase lag between the rhythms of photoreceptors’ sensitivity, assessed through pupillary light reflex, and of glare discomfort, suggests two independent underlying mechanisms. In conclusion, our findings highlight the need to take time-of-day and biological rhythmicity into account in the evaluation of light-induced discomfort glare. Apprehending these mechanisms may help understand photophobia in clinical populations, such as in migraine patients, and should be taken into account to optimize light quality at home and at the workplace, both for day and night work.

## Introduction

Discomfort glare, generally caused by an intense illumination, an excessive brightness or poorly positioned lights, is commonly experienced by healthy individuals. Photophobia, on the other hand, describes a sensory disturbance (generation or exacerbation of pain) provoked by an abnormal light sensitivity, and is a common symptom in neurological conditions such as migraine attacks (1, 2) or traumatic brain injury (1, 3).

Discomfort glare seems to be stronger in response to blue light, when compared to red light (4–6). These higher levels of discomfort glare induced by short-wavelength light exposures (4–6), are consistent with an involvement of intrinsically photoreceptive retinal ganglion cells (ipRGCs). Indeed, these photoreceptors, which are primarily implicated in non-visual functions (such as the photo entrainment of the biological clock or the pupillary light reflex (PLR) (7–13)), have a peak of sensitivity at 480 nm, and are therefore strongly activated by short wavelengths (blue light) (7–13). As ipRGCs project to multiple structures, including some regions involved in sensitivity and pain (14–16), they may play a key role in discomfort glare light sensitivity. The results regarding spectral sensitivity in photophobia is more controversial. Main and colleagues described higher discomfort under blue light exposure in between migraine attacks (17), but Chronicle et al. reported red light to be the least comfortable (18). The most recent studies on this topic revealed that white, blue, amber and red lights were all equally photophobic, with the exception of green light inducing less headache pain than the other light stimuli (19, 20), suggesting that photophobia may originate in cone driven retinal pathways. Altogether, these results reveal that the precise photobiological mechanisms underlying light-induced discomfort glare and photophobia remain poorly understood.

Time-of-day variations have been reported for certain sensory pathways. Fluctuations of olfaction and nociception across the 24-h day have been described, with a maximum olfactory sensitivity at 21:00 (21) and a peak of pain perception at 3:30 (22). These studies revealed a circadian modulation of sensory sensitivity, by the central circadian clock located in the suprachiasmatic nuclei (SCN) of the hypothalamus. Given that the circadian timekeeping system plays a key role in physiology, from gene expression to cortical activity and behavioral functions (23–28), it is likely to be involved in the regulation of all sensory perceptions, including visual perception and discomfort glare. However, to our knowledge, the effect of time-of-day on light-induced discomfort glare has never been systematically assessed.

Discomfort glare has mainly been evaluated subjectively. However, given that this response relies on physiological reactions, it may be assessed with objective measures such as pupil constriction. Indeed, although the relationship is not systematic (29), previous studies have reported correlations between pupil constriction and discomfort glare (30, 31), suggesting that the size of the pupil may be a correlate of discomfort glare. The magnitude of the pupillary light reflex (PLR) may also depend on the time of day. Two studies have revealed a 24-h variation of pupillary response to blue light, compatible with a diurnal variation of ipRGC sensitivity (32, 33). Zele and collaborators evaluated PLR sensitivity every hour in a 20-hour protocol and demonstrated a diurnal variation of the post-illumination pupil response (PIPR, a marker of the ipRGC response) in response to blue and red light, with a minimum light sensitivity around 23:00 (and no peak) (33). Münch and colleagues looked at the PIPR in response to red or blue light every hour during two 12-hour sessions (32) but solely identified a significant variation of the PIPR in response to blue light, with a minimum sensitivity to blue light around 7:30. A more recent study investigated cone sensitivity to light every 2 hours during a 24-h protocol by conducting a full field cone ERG and divulged both a linear decrease and a circadian rhythm, with a maximal response at 20:00 (34). Although these studies suggest that the circadian biological clock influences pupillary light sensitivity, no overall consensus on the timing of this rhythmicity has emerged and no circadian modulation of rods has ever been identified.

In this study, we aimed to determine whether light-induced discomfort glare displays rhythmicity over the 24-h day and to assess the precise contribution of the circadian system and sleep-related processes, by systematically assessing discomfort glare during a constant-routine protocol in highly controlled conditions. We also aimed to investigate the contribution of each photoreceptor through the assessment of the pupillary light reflex response as a proxy to their activity. We hypothesized that: 1) sensitivity to light would follow a circadian rhythm for both subjective discomfort glare and pupillary light reflex; 2) Discomfort glare would be correlated to photoreceptors sensitivity and predominantly driven by ipRGCs.

## Results

### Subjective evaluation of discomfort glare

Discomfort glare to light was assessed every two hours in response to red, orange and blue light spectra at three irradiances, throughout the whole 34-h constant routine protocol (Figure 5). The EC_50_ values (extracted from the Irradiance Response Curves [IRCs], see supplementary figure 4), which reflect the amount of light necessary to half-maximal discomfort glare (50% of the maximal response on the visual analog scale), were plotted as a function of time to observe the time-of-day effect (Figures 1A, 1D and 1G; all R² > 0.93). Discomfort glare to light across the 24h day was regressed by a linear and a sinusoidal model for each light spectrum (Figures 1A, 1B and 1C). Specifically, for red light, we found a linear component, showing that sensitivity to red light increases linearly with time spent awake (Figure 1D; R² = 0.87; p < 0.001), as well as a sinusoidal component, showing a circadian modulation of sensitivity to red light, with a peak of sensitivity at 2:30 (Figure 1G; R² = 0.88). For orange light, we reported a linear increase in discomfort glare with time spent awake (Figure 1E; R² = 0.98; p < 0.00001) and a sinusoidal fluctuation of sensitivity to orange light across the 24-h day with a maximum sensitivity at 3:30 (Figure 1H; R² = 0.98). For blue light, we identified a significant linear component, showing a small increase in discomfort glare with constant wakefulness (Figure 1F; R² = 0.44; p = 0.05), and a strong sinusoidal variation, revealing an important circadian modulation with a peak of sensitivity at 3:30 (Figure 1I; R² = 0.99).

**Figure 1.**
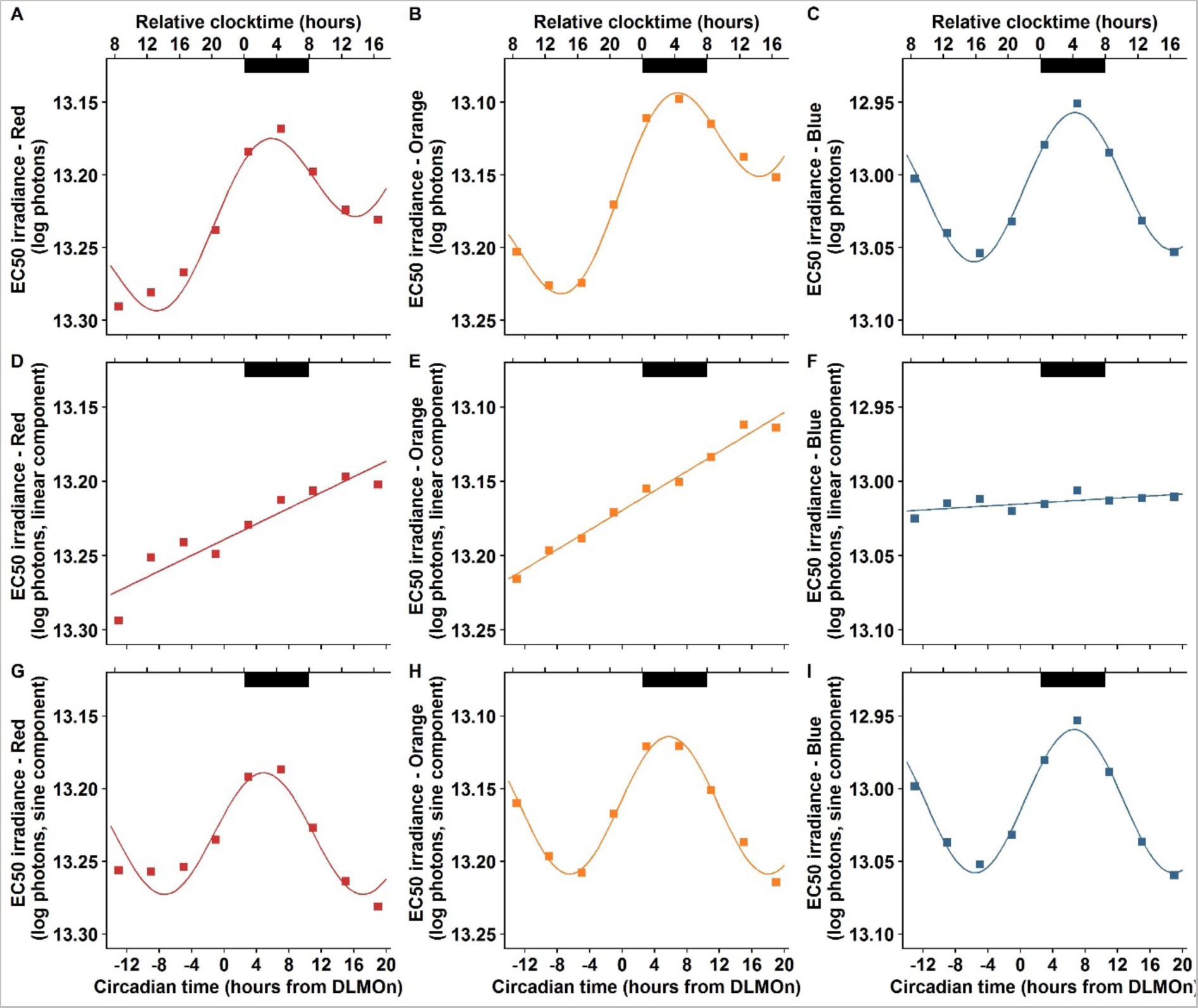
Mean discomfort glare in response to 10-second red (470 nm), orange (600 nm) and blue light (635 nm) exposures across the 34-h constant routine protocol (n = 12). Dark bars correspond to the average timing of habitual sleep episodes (biological night). Circadian time 0 corresponds to Dim Light Melatonin Onset (DLMOn, mean ≃ 21:30). A-C. Combined models (sum of linear and sinusoidal components) applied to raw EC_50_ values (extracted from irradiance response curves) for red light (A. R² = 0.93), orange light (B. R² = 0.99) and blue light (C. R² = 0.99). D-F. Linear components for red light (D. R² = 0.87; p < 0.001), orange light (E. R² = 0.98; p < 0.00001) and blue light (F. R² = 0.44; p = 0.052). Discomfort glare increases (EC_50_s decrease) with time spent awake for red and orange light. G-I. Sinusoidal components for red light (G. R² = 0.88), orange light (H. R² = 0.98) and blue light (I. R² = 0.99). Discomfort glare follows a circadian rhythm with a maximum of sensitivity at 2:30 (red light) or 3:30 (orange and blue light).

### Discomfort glare is predominantly ipRGC related

Altogether, these data show that subjective discomfort glare is rhythmic across the 24-h day, with a peak of sensitivity in the middle of the night (between 2:30 and 3:30; Figure 2A). As illustrated in Figure 2A, the overall irradiance necessary to induce moderate discomfort glare is always lower for blue light that for orange and red lights, at equal photon density. All three sinusoidal models are significantly different from each other (for all comparisons p < 4.9x10^-8^ after correction for repeated measures).

**Figure 2.**
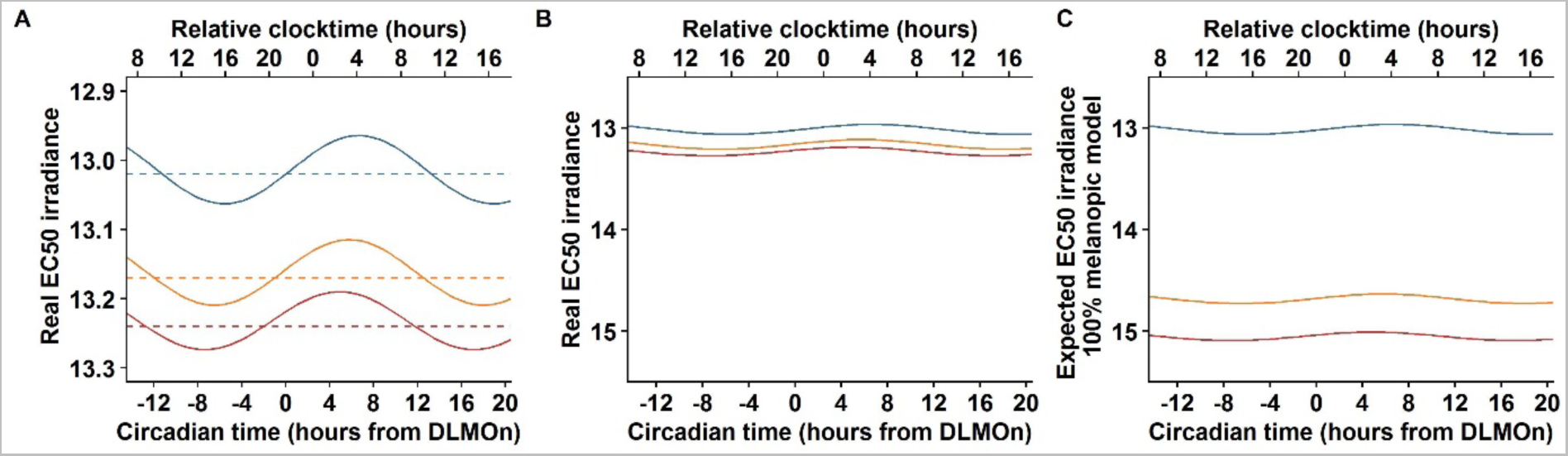
Circadian rhythms of discomfort glare in response to 10-second red (470 nm), orange (600 nm) and blue light (635 nm) exposures across the 34-h constant routine protocol (n = 12). Circadian time 0 corresponds to Dim Light Melatonin Onset (DLMOn, mean ≃ 21:30). EC50 irradiances are expressed in log photons. **A.** Real sinusoidal components for red (bottom red curve), orange (middle orange curve) and blue light (top blue curve). The dotted lines represent the mesor (mean sensitivity) of each rhythm: 13.24 log photons for red light, 13.17 log photos for orange light, and 13.02 log photons for blue light. All three sinusoidal models are significantly different from each other (all ps < 4.9 x 10^-8^). The peak of discomfort glare (minimum EC_50_ value) occurs at 2:30 for red light and at 3:30 for orange and blue light. **B.** Real sinusoidal components presented in **A.** rescaled in order to compare to the model exposed in **C.** For all three types of light, the mean sensitivity is close to 13 log photons. **C.** Expected sinusoidal components for red, orange and blue light, if discomfort glare was solely induced by activation of ipRGCs. Mean sensitivity for blue light is 13 log photons for blue light, 14.7 log photons for orange light and 15 log photons for blue light.

In order to further investigate how the different photoreceptors might contribute to discomfort glare, we extracted the melanopic content of each light stimulus, based on the ipRGC nomogram of sensitivity which peaks at 480 nm(35). As our orange light stimulus contained 50 times less melanopic energy than our blue light stimulus, the EC_50_ for orange light should be 50 times (1.7 log unit) higher than that to blue light, should discomfort glare be due to melanopsin only. Similarly, as our red light stimulus was a 100 times lower melanopic stimulus compared to blue light, the EC_50_ to red light should be 2 log units higher that to red light, again, if discomfort glare were to be only ipRGCs-related (Figure 2C). This means that if the EC_50_ with blue light is 13.02 log photons/cm²/s, then the expected sensitivity, based on the sole ipRGC input, should be 14.69 log photons/cm²/s with our orange stimulus and 15.06 log photons/cm²/s with our red stimulus (Figure 2C). Our results, illustrated in Figure 2B, are not those expected based on an involvement of melanopsin alone in discomfort glare (Figure 2C), and therefore allow to exclude the hypothesis that discomfort glare relies on a melanopsin-only model.

### Objective pupillary light reflex is rhythmic across the 24-h day

To further clarify the photobiological mechanisms involved in discomfort glare, we assessed the pupillary light reflex (PLR) in response to low intensity blue light, red light, and high intensity blue light, repeatedly over the 24 hours. The EC_50_ values, that were extracted from the IRCs of maximum pupil constriction in response to low intensity blue light and red light (18 curves; all R² > 0.93) were plotted as a function of time (Figures 3A and 3D; all R² > 0.85). Fluctuations of the post-illumination pupil response (6 s after the end of a high intensity blue light exposure, abbreviated as PIPR6s) were also observed on the combined model (sum of linear and sinusoidal components) (Figure 3G; R² = 0.95). We identified a linear decrease in pupil constriction (increase in EC_50_ and decrease in PIPR6s), as a function of time elapsed since waketime for low intensity (Figure 3D; R² = 0.27; p < 0.03) and high intensity blue light (Figure 3F; R² = 0.74; p < 0.00001). We did not find a linear trend for red light exposure (Figure 3E; R² = 0.13; p = 0.14). A significant sinusoidal rhythmicity was observed for all three types of light: low intensity blue light (Figure 3G; R² = 0.91), red light (Figure 1H; R² = 0.84) and high intensity blue light (Figure 3I; R² = 0.94). This circadian component revealed a peak sensitivity at 9:00 (low intensity blue light), 10:00 (red light) and 10:30 (high intensity blue light). Similar results were obtained for the maximum constriction in response to high intensity blue light (Supplementary figure 1). When PLR curves of low intensity blue light and red light are superimposed (Supplementary figure 2), these data show that although both circadian rhythms have a peak in the morning (9:00 – 10:00), blue light induces more pupil constriction than red light, at equal photon density (p < 2.2 x 10^-16^).

**Figure 3.**
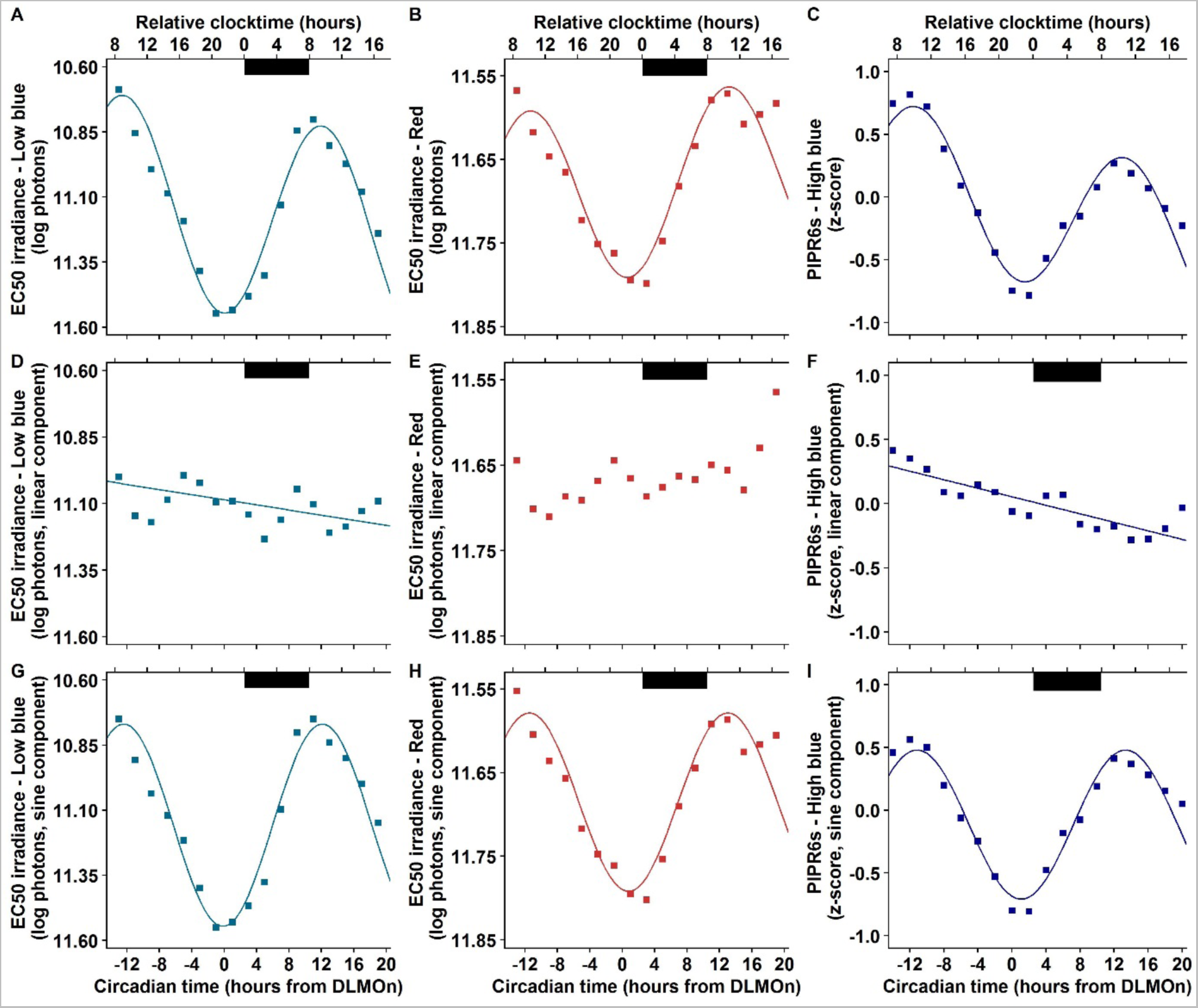
Mean pupillary light response (PLR) to 1-second low intensity blue (465 nm), red (630 nm) and high intensity blue light (465 nm) exposures across the 34-h constant routine protocol (n = 12). Dark bars correspond to the average timing of habitual sleep episodes (biological night). Circadian time 0 corresponds to Dim Light Melatonin Onset (DLMOn, mean ≃ 21:30). **A-C.** Combined models (sum of linear and sinusoidal components) applied to raw EC50 values for low intensity blue light (**A.** R² = 0.92), red light (**B.** R² = 0.85) and applied to the post-illumination pupil response 6s after high intensity blue light exposure (PIPR6s) (**C.** R² = 0.95). **D-F.** Linear components for low intensity blue light (**D.** R² = 0.27; p < 0.03), red light (**E.** R² = 0.13; p = 0.14) and high intensity blue light (**F.** R² = 0.74; p < 0.00001). PLR decreases (EC_50_ increase and PIPR6s decrease) with time spent awake for low and high intensity blue light. **G-I.** Sinusoidal components for low intensity blue light (**G.** R² = 0.91), red light (**H.** R² = 0.84) and high intensity blue light (**I.** R² = 0.94). PLR follows a circadian rhythm with a maximum of sensitivity at 9:00 (low intensity blue light), 10:00 (red light) or 10:30 (high intensity blue light).

### Discomfort glare is not explained by pupil constriction, nor by baseline pupil diameter

The temporal phase relationship reveals that the peak of pupillary light reflex does not align with the peak of discomfort glare, but that there is a 6.5-hour phase lag between the two peaks (Figure 4A). The circadian rhythmicity of discomfort glare and PLR are not explained by the rhythmicity of the baseline pupil size, as temporal phase lags are of 5-h for discomfort glare and 11.5-h for PLR (Figures 4B and 4C).

**Figure 4:**
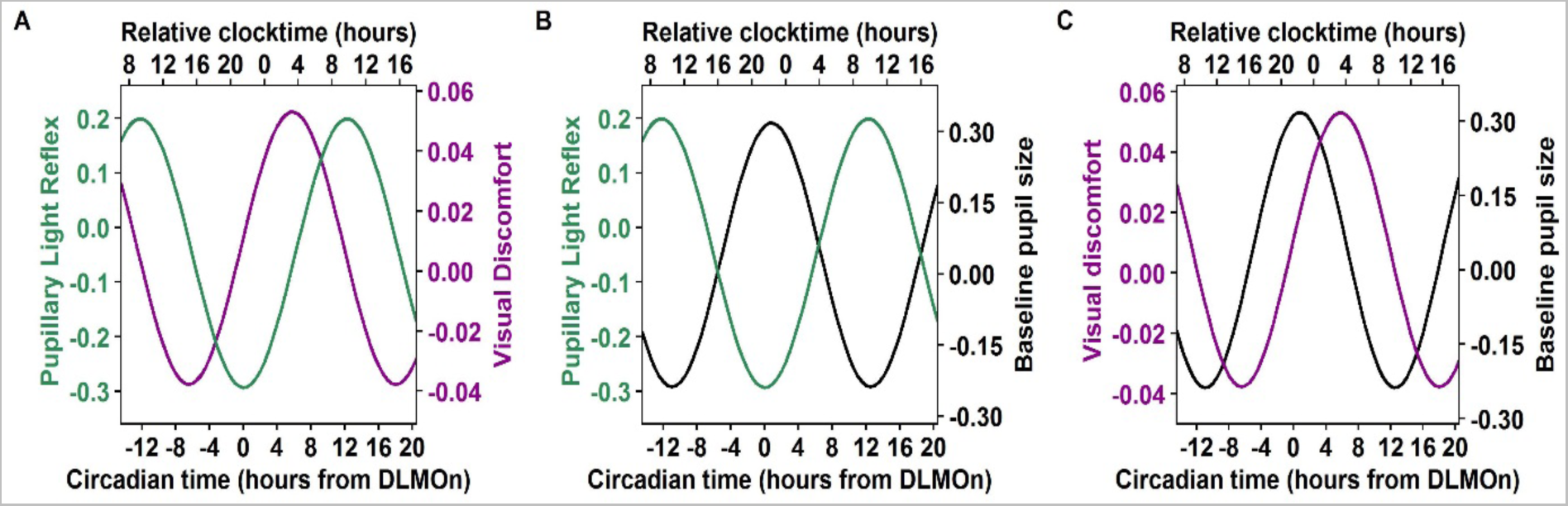
Phase relationships between circadian components of visual discomfort, pupillary light reflex and baseline pupil diameter across the 34-hour constant routine protocol (n = 12). Circadian rhythm of baseline pupil diameter data was taken from Daguet et al. (2019). Circadian time 0 corresponds to DLMOn (mean DLMOn = 21:30). **A.** Circadian rhythms of discomfort glare (purple curve; peak at 3:30) and pupillary light reflex (green curve; peak at 10:00). **B.** Circadian rhythms of pupillary light reflex (green curve; peak at 10:00) and baseline pupil size (black curve; peak at 22:30). **C.** Circadian rhythms of discomfort glare (purple curve; peak at 3:30) and baseline pupil size (black curve; peak at 22:30).

### Light sensitivity variations across the 24-hour day are mainly due to the circadian timing system

We investigated the relative contributions of sleep and circadian drives to light sensitivity, by calculating the mean changes in both these components and expressing them relatively to the total amplitude over 24 hours. Both for discomfort glare and pupillary light reflex, we found that the time-of-day effect is mainly related to the circadian system rather than to the sleep/wake cycle. The circadian system accounted for approximately 80 % of the full magnitude of discomfort glare changes over 24 hours (71 %, 68 %, 96 % for red, orange and blue light respectively), the remaining 20 % being accounted for by the homeostatic component. The relative contributions of sleep and circadian processes for discomfort glare in response to blue light are illustrated in figure 5. For the pupillary light reflex, the circadian clock was responsible for 90% of the time-of-day effect (93 %, 94 % and 85 % for low intensity blue light, red light and high intensity blue light respectively), revealing that only 10 % is sleep-related.

**Figure 5:**
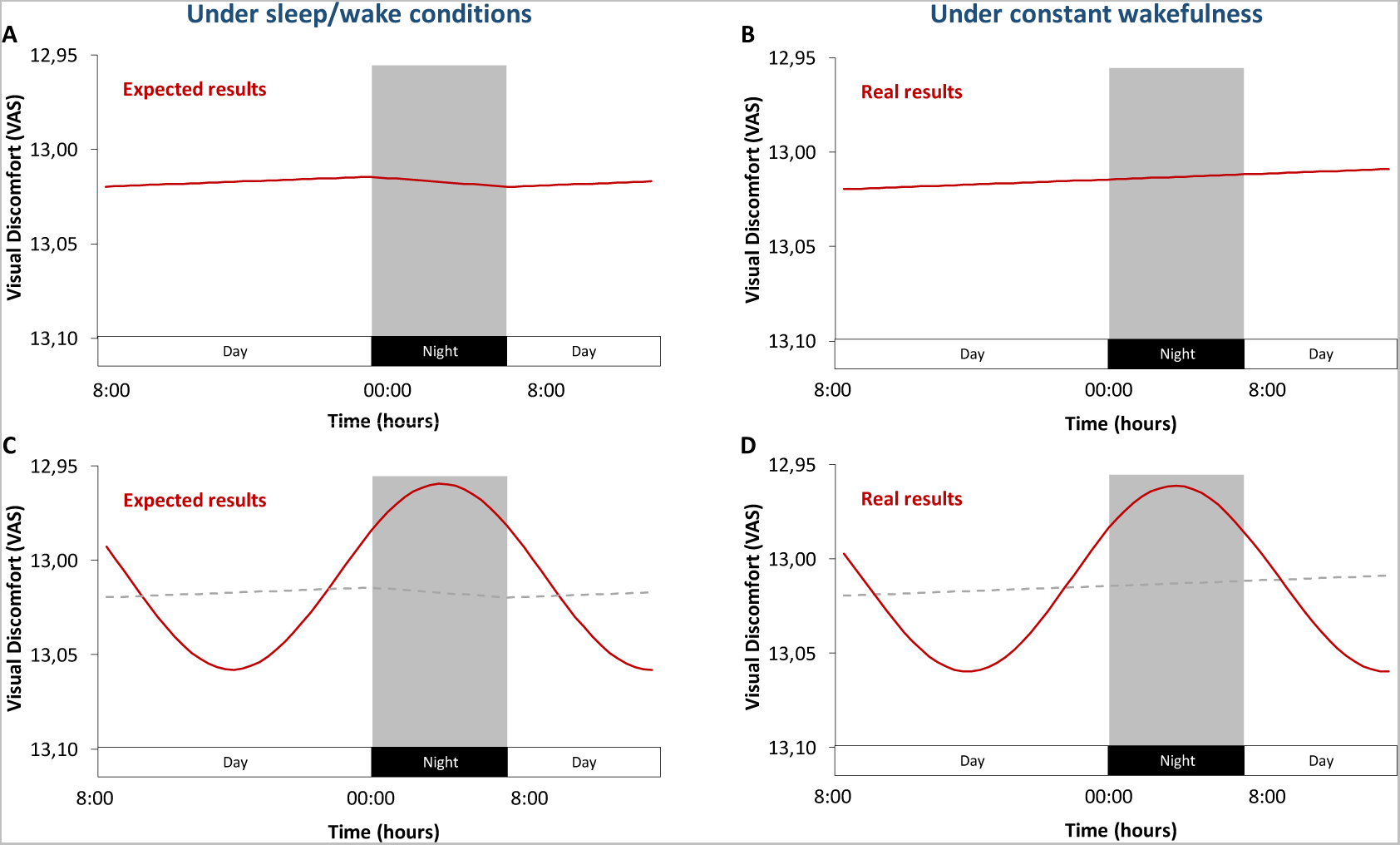
Variations of discomfort glare in response to blue light across the 24h day. **A-B.** Expected discomfort glare, according to the current homeostatic model. Under regular sleep/wake conditions (**A**), discomfort glare increases during wakefulness, in parallel to sleep pressure, and decreases during the night. Under constant wakefulness conditions (**B**), discomfort glare increases during wakefulness, and keeps increasing in the absence of sleep. In this model, discomfort glare only depends on time since awakening (during the day), and time since bedtime (at night). **C-D.** Our results show that discomfort glare is driven by two independent and additive components: a homeostatic drive and a circadian drive. With sleep at night (**C**) both mechanisms co-exist. Discomfort glare oscillates sinusoidally (circadian drive) and increases linearly during wakefulness and decreases during sleep (homeostatic drive – grey dotted line). Without sleep at night (**D),** our results under constant wakefulness show the superimposed additive homeostatic (grey dotted line) and circadian regulation of discomfort glare, with both a linear increase with time spent awake, and a sinusoidal oscillation.

## Discussion

Our results demonstrate that discomfort glare in response to light exposure is rhythmic over the 24-hour day and results from two additive components. They reveal that discomfort glare is strongly driven by the circadian timing system (≃ 80%) and oscillates sinusoidally over the 24 hours, with light sensitivity maximal in the middle of the night (at 3:30), and minimal during the afternoon (at 15:30). They also show that discomfort glare increases linearly during prolonged wakefulness, possibly driven by a sleep-related mechanism (≃ 20%). Our results also show that the light sensitivity of all three types of photoreceptors (rods, cones and ipRGCs) evolves circadianly over time, but does not oscillate in phase with the rhythm of discomfort glare. The peak of non-visual sensitivity to light (pupillary light reflex) occurs in the morning (at 10:00), which suggests that the pupil response is neither a proxy for, nor involved in, discomfort glare. Finally, our results demonstrate that discomfort glare is primarily driven by ipRGCs, and that rods and cones are also involved in its regulation.

### Discomfort glare is rhythmic and mainly but not exclusively ipRGC related

As no previous studies had evaluated the 24-h rhythm of discomfort glare, this is the first evidence that visual sensitivity is rhythmic and results from the additive input of two regulatory components: the circadian timing system and the homeostatic sleep-related mechanism. Although the precise physiological role of discomfort glare is unclear, one view suggests that discomfort glare (which could be viewed as a light-induced pain) is a protective response that discourages us from viewing intensely bright objects that could harm the retina. The purpose of a 24-h rhythmicity of discomfort glare (involving homeostatic and circadian processes), remains unclear, but shares common pathways with pain sensitivity which also peaks in the middle of the night (22). The linear increase in discomfort glare with sleep debt (homeostatic process) is also in line with the previously reported increase in sensory sensitivity (pain (22), audition (21)) and visual attention (36) with time elapsed since waketime.

Independently of the time-of-day effect, our results show a wavelength specific effect which reveals that healthy individuals report higher levels of discomfort glare after exposure to blue light than after orange and red light exposure (as the light intensity necessary to induce a moderate subjective sensation of discomfort glare is the lowest for blue light). As this narrowband blue light (peaking at 470 nm) predominantly activates ipRGCs, this result suggests a major role of ipRGCs in discomfort glare. Previous studies had suggested a role of ipRGCs in discomfort glare both in healthy individuals (6) and migraine patients (14). Precisely, Noseda and colleagues reported a prevalent exacerbation of migraine by light exposure among blind individuals who had a massive rod/cone degeneration, therefore suggesting an implication of ipRGCs (14). Our analysis confirms the implication of ipRGCs, but also reveals that they are not solely responsible for discomfort glare. Precisely, as the levels of discomfort glare observed under red and orange light conditions (targeting more cones that ipRGCs) are only modestly different in terms of EC_50_ values, with a difference of less than 0.22 log photons and not of 1.7 log units as predicted by the sole ipRGC model, we argue that rods and/or cones also play a large role in discomfort glare. S-cones (peaking at 440 nm) had previously been suggested to be involved in discomfort glare (37), but a recent study reported that the specific activation S-cones did not strongly affect visual comfort, glare or brightness (38). Despite obvious differences between healthy individuals and migraine patients, our data on the relative contribution of each photoreceptor in discomfort glare may explain why Noseda and colleagues published contradictory results on the role of ipRGCs (14), cones (20) and rods (39) in migraine photophobia and did not reach a general consensus regarding the photobiological mechanisms involved in discomfort glare (or photophobia). However, the involvement of the ipRGC pathway in the transduction of light information into a painful perception has been put forward due to the projection of ipRGCs directly to pain centers in the posterior thalamus (14–16). This connection may be a significant part of the “photophobia pathway”, which could be pathologically increased in neurological conditions (40). The 24-h rhythmicity of discomfort glare reported in this study, could reflect the activity of this circuitry.

### Pupillary light reflex and photoreceptor sensitivity is rhythmic

Though it is difficult to isolate the function of each photoreceptor in the different phases of the PLR, studies have suggested that rods and cones primarily contribute to the phasic response of the PLR (maximum constriction) whereas melanopsin is mainly involved in the steady and tonic phases (PIPR) (41–45). Specifically, authors agree that ipRGCs are involved in pupil constriction in response to light, but S-cones (46), M-cones and L-cones (47) have been proposed to have an inhibitory effect on the pupil constriction, or even a positive effect (44). Our results reveal that both metrics of the pupillary light reflex (maximum constriction as a marker of rod and cone input, and PIPR6s as an index of ipRGC input) are also strongly regulated by the circadian clock, with a peak of sensitivity in the morning (9:00 - 10:30). This suggests a circadian modulation of the sensitivity of all photoreceptors: rod, cone and ipRGCs, either locally at the retinal level or centrally through a downstream structure. Within this time-of-day effect, the linear decrease in photoreceptor sensitivity (decrease in PLR response) with sleep debt is not surprising, as the sensitivity of certain non-visual functions, such as the circadian phase shifts in response to bright light, also decreases with sleep deprivation(48, 49). These results are also in line with results from Zele and collaborators, who identified a rhythmic PIPR to both blue and red light (488 nm and 610 nm respectively), although with a peak at a different phase, in the late evening (33). Our results, however, differ from the results by Münch and colleagues showing a 24-h variation of PIPR following blue light exposure, with a maximal response around 7:30, but no time-of-day effect in response to red light, suggesting no circadian variation of cones’ sensitivity (32).

### Discomfort glare does not rely on pupil constriction

As discomfort glare and pupillary light reflex are both controlled by the circadian clock, it is legitimate to think that variations in the pupil response could explain the changes in discomfort glare. In this direction, correlations between pupil diameter and subjective discomfort glare have been reported (30, 31). However, the 6.5-hour phase lag between the peak of pupillary light reflex (at 10:00) and the peak of discomfort glare (at 3:30) in our data shows that the two responses do not covary and suggests that these responses rely on two independent mechanisms. Indeed, the hypothesis that the peak of discomfort glare is caused by a saturation effect (the pupil has reached its maximum of constriction and is unable to constrict more), is invalidated by the phase relationships we found. In other words, the circadian rhythmicity of discomfort glare is not explained by the circadian rhythmicity of the photoreceptors’ sensitivity, therefore suggesting that it may not be regulated at the retinal level.

In terms of photobiological mechanisms, the visual pathway consists in the transmission of the light signal from the rods and cones through the retinal ganglion cells, which relays in the lateral geniculate nucleus or the superior colliculus and finally reaches the visual cortex (50). On the other hand, the non-visual pathway starts at the ipRGC level and, through the retino-hypothalamic tract, activates a number of brain structures such as the ventrolateral preoptic nucleus (VLPO), the locus coeruleus (LC) and the olivary pretectal nucleus (OPN) (15). The discomfort glare pathway is less clear, but is likely to involve both cortical and subcortical cognitive and emotional structures, such as the thalamus or the amygdala (51–53). Knowing that homeostatic and circadian modulations of cortical activity (54) and excitability (27) have been previously identified, discomfort glare may be regulated at the neuronal level. More precisely, functional MRI results from Vimal and collaborators (2009) (55) suggest that the time-of-day changes in discomfort glare come from an increased SCN responsiveness and not from higher activation in the visual cortex.

### Baseline pupil diameter does not explain discomfort glare, nor pupillary light reflex

Independently from the pupillary light reflex, another hypothesis could be that the intensity of discomfort glare depends on the amount of light entering the eye, itself relying on the dilation of the pupil. Yet, this hypothesis seems unlikely as the peak of discomfort glare (at 3:30) does not coincide with the time at which the pupil diameter is maximal (22:30). This result supports the idea that the pupil diameter by itself cannot be used as an objective indicator of the degree of glare (Hopkinson, 1956). Similarly, a 11.5h phase lag is observed between the peak of pupillary light reflex (at 10:00) and the time at which the pupil diameter is maximal (22:30), again suggesting that these responses involve two distinct pathways.

Light exposure activates the parasympathetic pathway, and inhibits the sympathetic pathway, therefore leading to pupil constriction (44, 56–58). One hypothesis is that the circadian modulation of the pupil response that we observed involves changes in the autonomic nervous system (57) and that the maximal PLR in the morning (at 10:00) is explained by a higher activity of the sympathetic pathway, and a lower parasympathetic activity.

## Limitations

Our study has a few potential limitations: First, the participants included in this study were only men. Sensitivity to light has been shown to differ in men and women, with women showing lower brightness perception than men (59), as well as higher relative pupil constriction amplitudes (60). Nonetheless, it has been repeatedly shown that circadian physiology is similar in men and women, with only minor differences such as a slightly larger amplitude(61, 62), and a slightly shorter circadian period in women (63). Second, the pupillary light reflex and discomfort glare were not measured at the same time, but with a delay of about 20 minutes between them. Given that we are addressing circadian rhythmicities and not ultradian frequencies here, and given the 6.5 h delay we found between rhythmicities of discomfort glare and of pupillary response, we exclude that the small delay between the 2 measures could be an issue. Third, our protocol enabled us to target photoreceptors with only a relative specificity.

The adequate approach to precisely separate each photoreceptor contribution would make use of the silent substitution paradigm (metameric lights) (64). However, the time required to properly separate each photoreceptors’ contribution would not allow measures as frequent as ours in this study (every 2 hours). In addition, although this would clarify the photoreceptor contribution, we believe it is unlikely that it would change the relative contribution and the circadian phase (timing) of the non-visual sensitivities we found. Therefore, our conclusion that circadian rhythmicity of the photoreceptor’s sensitivity and pupil diameter does not drive circadian rhythmicity of discomfort glare would remain unchanged.

## Conclusion

In conclusion, our results show that discomfort glare in response to red, orange and blue light is rhythmic over the 24-hour day and controlled by two superimposed processes: a strong circadian drive and a modest homeostatic sleep-related component. The ipRGC pathway is the primary transducer of discomfort glare, but cones seem to play a relatively large part in the response as well. The 6.5-hour phase-lag we found between the peak of discomfort glare and the peak of pupillary light reflex suggests two independent underlying mechanisms and demonstrates that pupillary constriction is neither a proxy for, nor involved in discomfort glare. Further studies are required to decipher the mechanisms at the origin of discomfort glare and to determine the precise role of each photoreceptor.

## Material & methods

### Participants

Twelve healthy men (20 - 29 years old, mean age = 22.7 ± 3.3 years; BMI = 21.8 ± 3.1 kg/m²) were included in this study. Neurological, psychiatric and sleep disorders were excluded by clinical examination and psychological questionnaires (Pittsburg Sleep Quality Index Questionnaire and Beck Depression Inventory)(65, 66). Participants had an intermediate chronotype (Horne and Ostberg Chronotype Questionnaire score between 31-69) (67) and had not done any shift work, or experienced transmeridian travel during the previous three months. Participants had normal visual acuity (Landolt Ring Test and Monoyer scale), contrast vision (Functional Acuity Contrast Test) and color vision (Farnworth D-15 and Ishihara Color Test). All experimental procedures were carried out in accordance with the Declaration of Helsinki. The study was approved by the local research ethics committee (CPP Lyon Sud-Est II) and participants provided written informed consent for participation.

### Study Design

Participants were asked to maintain a regular sleep/wake schedule (bedtimes and waketimes within ± 30 minutes of self-targeted times) for an average of three weeks before admission to the laboratory, with verification by wrist activity and light exposure recordings (ActTrust, Condor Instruments, São Paulo, Brazil). Subjects were then admitted to the laboratory for a 56-hour experimental protocol (Figure 5), in which they were kept in an environment free from external time cues (clocks, television, smartphones, internet, visitors, sunlight etc.). Subjects maintained contact with staff members specifically trained to avoid communicating time-of-day information or the nature of the experimental conditions to the subjects. Participants arrived at about 10:00 on the first day. They were allowed to familiarize themselves with the laboratory environment, low light levels (< 0.5 lux), equipment, and measurements. Lunch and dinner were served at about 12:30 and 19:00. A series of measurements were then performed until bedtime (participant’s habitual bedtime), and an 8-hour sleep episode was scheduled (constant darkness; recumbent position). This was followed by a 34-hour constant-routine protocol beginning at the participant’s usual waketime on day 2, and ending on day 3 (18:00 on average). Habitual bedtimes were determined on the basis of sleep times averaged over the seven days preceding the laboratory segment of the protocol. Average bedtime was 23:45 and average waketime was 8:00.

### Constant Routine Protocol

A constant routine (CR) paradigm was used to reveal the endogenous circadian rhythmicity of various parameters. The CR was conducted under constant environmental conditions, to eliminate, or distribute across the circadian cycle, the physiological responses evoked by environmental or behavioral stimuli (i.e. sleeping, eating, changes in posture, light intensity variations) (68, 69). In practical terms, participants were asked to remain awake for 34 hours (starting at their habitual waketime), with minimal physical activity, while lying in a semi-recumbent (45°) posture in bed. This posture was also maintained for the collection of urine samples and bowel movements. Room temperature (mean = 23 °C ± 0.6 (SD)) and ambient very dim halogen light levels were kept constant. Light intensity was homogeneous in the room (< 0.5 lux at the participant’s eye level in all directions of gaze). Participants were given small equicaloric snacks and fluids at hourly intervals, to maintain an equal nutritional caloric intake and stable hydration over the circadian cycle. Caloric requirements were calculated on the basis of basal metabolic rate determined with the Wilmore nomogram and were adjusted upward by a 7 % activity factor (70, 71). Fluid intake was calculated for each subject, to account for the sedentary nature of the CR(71). A member of the study staff remained in the room with the participant at all times during the CR, to monitor wakefulness and to ensure compliance with the study procedures.

**Figure 6:**
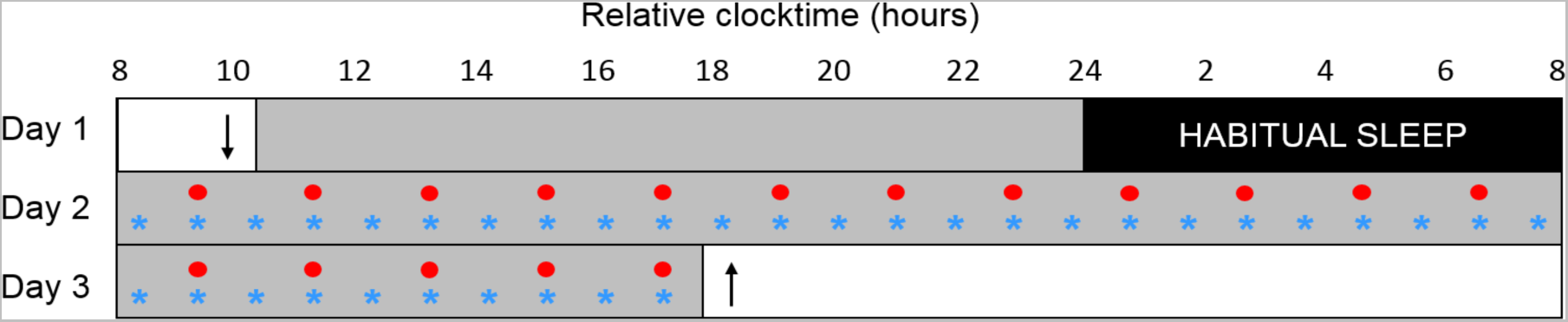
Overview of the experimental protocol. After a day of habituation (day 1) and an 8-h sleep episode, participants were subjected to a 34-hour constant routine (CR: days 2 and 3). Melatonin levels were assessed hourly (blue stars); visual discomfort and pupillary light reflex were evaluated every two hours (red circles). Participants arrived at about 10:00 on day 1 (down arrow) and left the laboratory at about 18:00 on day 3 (up arrow). Gray rectangles represent wakefulness in dim light (∼ 0.5 lux) and black rectangles represent scheduled sleep in darkness.

### Discomfort glare

Discomfort glare was assessed in response to light exposures at 3 different wavelengths: 470 nm (blue light), 600 nm (orange light) and 635 nm (blue light) (Supplementary figure 3) (Ledmotive, Barcelona, Spain). The lights at different wavelengths allowed us to activate relatively selectively different photoreceptors and therefore investigate the photobiological mechanisms involved in discomfort glare. Indeed, the narrowband blue light at 470 nm predominantly activates ipRGCs, large spectrum orange light (600 nm) activates predominantly M- and L-cones (but also affects, to a lower extent, rods and ipRGCs) and the narrowband red light at 635 nm majorly activates M-cones and L-cones (Supplementary table 1). For each wavelength, participants were exposed to 3 rising irradiances and light spectra were calibrated to provide the following corneal photon flux : 7. x *1012 photons/cm²/s (12.85 log photons); 1.3 x 1013 photons/cm²/s (13.11 log photons) and 2.5 x 1013 photons/cm²/s (13.4 log photons).

Light was measured at the eye level of participants in the experimental situation with a spectroradiometer (S-BTS2048, GigaHertzOptik, Germany) in order to determine spectra and irradiances. Photopic, melanopic and other alpha-opic illuminances (35) are given in Supplementary Table 1.

Background light intensity was low (< 0.5 lux) and homogeneous in the room. For each stimulus, the light was turned on, remained at the target irradiance for a duration of 10 seconds, and was then turned off. Participants were asked to keep their eyes open (attempting not to blink) and to look straight ahead directly at the light source (located at 150 centimeters from the participant’s eye; angle of the light spot ≃ 0.5°) during the totality of the light exposure (for setup illustration see Supplementary figure 4). Immediately after lights off, the discomfort glare, induced by each of the 9 light exposures, was assessed with a 100-mm computerized visual analog scale (VAS). Participants were asked to rate the intensity of the discomfort glare on the VAS, extending from “no visual discomfort”» (minimal score of 0/10) to “maximum imaginable visual discomfort” (maximal score of 10/10) (for similar procedures see (31, 72)). Stimuli were separated by an interval of at least 30 s of darkness.

As discomfort glare was measured at three arbitrary irradiances, we used a sigmoidal modelling approach (Supplementary figure 4) to extract the overall light sensitivity value (EC_50_) for each light spectra: red, orange and blue light. This is a better approach to the assessment of sensitivity, as it can be used to determine the half maximal effective irradiance, or EC_50_, corresponding to the irradiance required to induce 50% of the maximal response. The data were modeled with a sigmoidal function:

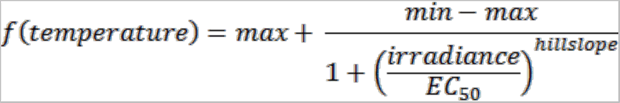

As the VAS is a bounded scale, minimum (min) and maximum (max) discomfort glare scores were set at 0 and 10, respectively. Hillslope, the slope of the curve, and EC_50_ were left free. The statistical power of the modeling approach was increased by calculating sigmoidal fits over 4-hour time epochs, corresponding to 2 evaluations of discomfort glare to each of the 3 irradiances (12.85, 13.11 and 13.4 log photons), providing six points on the regression curve (Supplementary figure 4). The EC_50_ values were extracted from each of the nine sigmoidal regressions (see formula above and Supplementary figure 4) and plotted as a function of time to observe the time-of-day effect (Figure 1).

In order to further investigate how the different photoreceptors contribute to discomfort glare, we calculated the melanopic content of each light stimulus, based on the ipRGC nomogram of sensitivity which peaks at 480 nm (supplementary figure 1)(35, 73). As our orange light stimulus contained 50 times less melanopic irradiance than blue light, 50 times (1.7 log unit) more light should be necessary to induce the same amount of discomfort glare. Similarly, as red light contains 100 times less melanopic content than blue light, the average sensitivity (EC_50_) irradiance should be 2 log units higher. These theoretical values, were then compared to the real data obtained.

### Pupillary Light Reflex

Pupillary light reflex was recorded every 2h with a hand-held monocular video-pupilometer (Neurolight, IDMed, Marseille, France). This device, placed at 25 mm from the cornea surface, detected pupil margins under infrared illumination (two infrared LED lights with a peak at 880 nm) and continuously tracked the pupil diameter. The pupilometer was placed in front of the participant’s left eye and held steadily by the experimenter. The participant was asked to keep the left eye wide open (without blinking) and to look straight ahead. During the measurement, the experimenter could see the pupil on the screen of the device and check that the device was correctly placed on the participant’s eye. This measurement was conducted in complete darkness as one eyelid was closed and covered by the participant’s hand and the other eye was covered by the device. Before each measurement, we also questioned the participant in order to ensure that the participant did not detect any ambient lighting. Pupil diameter was recorded with a sampling rate of 62 Hz and stored in mm in the output file of the pupilometer. Pupil diameter was considered abnormal when values were above 9 mm or below 2 mm. Artefacts were defined when an absolute change between 2 samples (sampling rate of 62Hz) was above 0.15mm (which corresponds to a change of approximately 9.3 mm per second). The baseline pupil diameter was detected over a 5 s segment in darkness and the PLR was measured in response to 1-second light exposures at 2 wavelengths and different irradiances in order to target relatively specifically each photoreceptor: rods (low intensity blue light: 1.17 x 10^9^, 3.5 x 10^9^, 1.17 x 10^10^, 3.5 x 10^10^ and 1.17 x 10^11^ photons/cm²/s at 465 nm), cones (red light: 1.6 x 10^11^, 4.8 x 10^11^, 1.6 x 10^12^, 4.8 x 10^12^ and 1.6 x 10^13^ photons/cm²/s at 630 nm) and ipRGCs (high intensity blue light: 10^14^ photons/cm²/s at 465nm) (see the blue and red light spectra in Supplementary figure 5).

Two parameters were analyzed: maximum constriction (in response to low intensity blue light at 465 nm and in response to red light at 630 nm to target M-cones and L-cones) and post-illumination pupil response at 6 seconds after lights off (PIPR6s; in response to high intensity blue light at 465 nm to target ipRGCs).

Maximum pupillary constriction and PIPR were calculated as a percentage of change relative to baseline levels of pupillary constriction, using the following equations:

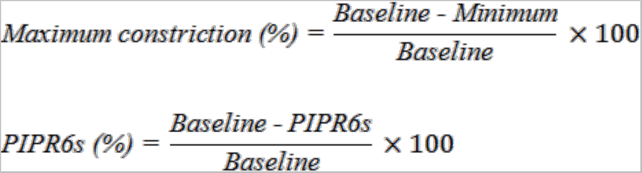

For a more precise assessment of the pupil responses (than what could be achieved with using the individual responses to the different light irradiances), intensity response curves were calculated for low intensity blue light and red light stimulations (as we did for discomfort glare – see supplementary figure 4). This is a better approach to the assessment of sensitivity, as it can be used to determine the half maximal effective irradiance, or EC_50_, corresponding to the irradiance required to induce 50% of the maximal pupillary response and reflects the sensitivity of the photoreceptors. The data were modeled with a sigmoidal function:

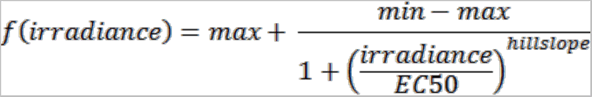

As previous studies had identified a maximal pupil constriction As previous studies had identified a maximal pupil constriction ranging from 60-65% (42, 74) and 80% (75), we chose to set the maximum (max) pupil constriction at 75%. The minimum (min) was set at 0, as no pupil constriction is observed in darkness. Hillslope, the slope of the curve, and EC_50_ were left free. Sigmoidal fits were calculated over 2-hour time epochs, corresponding to the evaluation of PLR at five irradiances, providing five points on the regression curve. The EC_50_ values were extracted from each of the eighteen sigmoidal regressions (see formula above) and plotted over time (Figure 3).

### Melatonin

Saliva was collected hourly, with cotton swabs placed directly in the mouth of the participant (Salivettes, Sarstedt, Nümbrecht, Germany). Samples were stored at -20 °C until centrifugation and assay. Melatonin levels were measured with an in-house radioimmunoassay ^125^I (RIA). This assay was based on a competition technique. The radioactive signal, reflecting the amount of ^125^I-labeled melatonin, was therefore inversely proportional to the concentration of melatonin in the sample. The sensitivity of the assay was 1.5 pg/mL. The inter-assay coefficients of variation for high (18.5 pg/mL) and low (10 pg/mL) melatonin-concentration controls were 19% and 22% respectively, and the mean intra-assay coefficient of variation was below 10 %. We determined the circadian melatonin profile of each participant over a 24-hour day, by applying a three-harmonic regression individually to the raw data collected during the CR (days 2 and 3)(76). The model equation was:

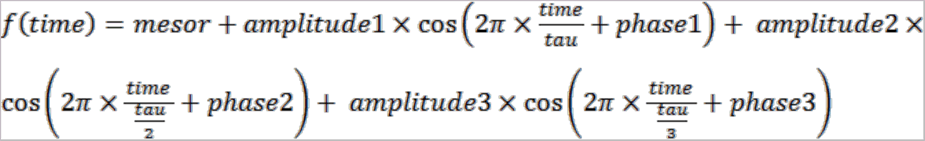

In the model, Tau (the circadian period) was constrained between 23.5 and 24.5 h; mesor, amplitudes (1 to 3) and phases (1 to 3) were set free.

The dim light melatonin onset (DLMOn), corresponding to the circadian phase, was calculated for each participant. DLMOn was defined as the time at which the ascending phase of the melatonin profile crossed the 25 % threshold of the peak-to-trough amplitude of the fitted curve. Due to technical problems with some saliva samples, the full 24-hour melatonin profile could not be obtained for two participants. For one of these participants, DLMOn was calculated on the basis of melatonin levels during the habituation day (day 1), rather than during the CR, for which we could not determine melatonin concentrations. For the second participant, in the absence of melatonin concentration data (flat profile below the limit of quantification of the assay), DLMOn was estimated from the mean phase angle calculated between habitual bedtime and DLMOn (based on (76)).

### Data Analysis

Outliers were identified on the basis of normalized data (*z*-scores) and were excluded from subsequent analyses (outlier.test, R, Version 3.6.1 - 2019-07-05, R Foundation for Statistical Computing, Vienna, Austria). We reduced inter-individual variability, by normalizing all data (except melatonin concentrations) by calculating individual *z*-scores and smoothing them with a weighted moving average (calculated on 3 points; 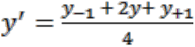). In order not to lose the first and last points of the times series, we calculated a truncated moving average on those 2 points (first point: 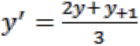; last point: 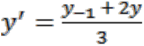).

The endogenous circadian phase was taken into account for each participant, by aligning the data with the onset of melatonin secretion (DLMOn). As DLMOn occurred at different times in different participants, individual melatonin onset values were set to 0 (DLMOn = circadian time 0), and all measurement times are expressed relative to melatonin onset. We modeled the effects of time on the responses observed during the 34-hour constant routine, using an additive model including a linear component (homeostatic, process S) and a sinusoidal component (circadian, process C). The equation of the combined model was:

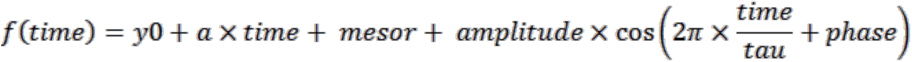

Tau (circadian period) was constrained between 23.5 and 24.5 hours (63, 77), whereas all other parameters were left free. Once the parameters of the combined model had been defined, process S and process C were modeled separately. The homeostatic component (process S) was regressed against the linear component of the model: *f*(*time*) = *y*0 + *α* × *time*. The circadian rhythmicity (process C) of the data was regressed against the sinusoidal component of the model: 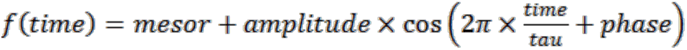.

A non-linear multiple regression approach followed by an anova, on individual and pooled datasets, was used to compare the sinusoidal models in response to red, orange and blue light for discomfort glare and in response to red and blue light for pupillary light reflex (78). As three comparisons were conducted, a multiple comparisons Bonferroni correction was applied to the p-values.

In order to compare the previously published data (25) to light sensitivity data presented here, baseline pupil diameter data were realigned according to each individual’s internal circadian time. Statistics were calculated with R (Version 3.6.1 - 2019-07-05, R Foundation for Statistical Computing, Vienna, Austria). Results were considered significant if *p* ≤ 0.05.

## Acknowledgments

We wish to thank all the volunteers who participated in this study. We also wish to thank the staff and students, and especially Pauline Kirchhoff who participated in data collection and analysis. A special thanks also goes to Dr Alain Nicolas who conducted the medical and physical examinations, and Dr Abhishek Prayag who shared with us his knowledge in mathematical modelling with R.

## Data Availability

The data that support the findings of this study are available from the corresponding author upon reasonable request.

## Author contributions

The experiment was conceived by CG, and designed by CG and ID. Data collection was performed by ID and CG. Data analyses were conducted by ID and CG, and hormonal assays were conducted by VR. ID, VR and CG interpreted the data. ID and CG wrote the manuscript, and VR edited the manuscript. All authors agreed to be accountable for all aspects of the work.

## Supplementary information

**Supplementary figure 1:**
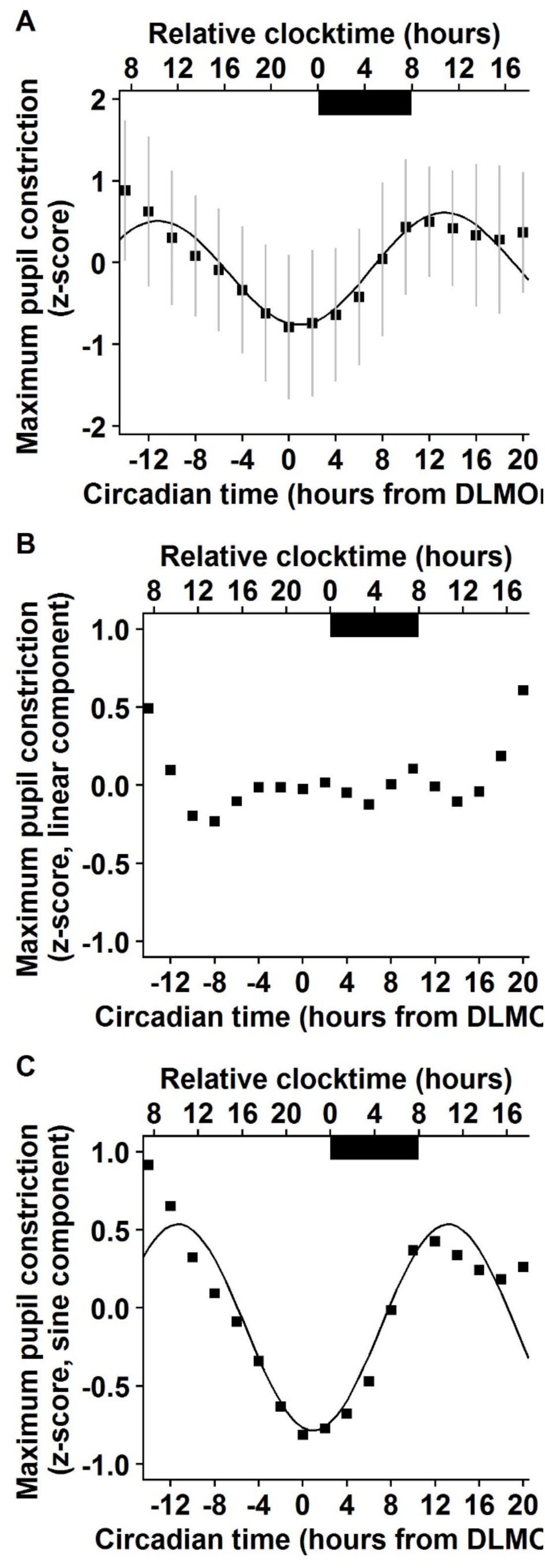
Mean maximum pupil constriction in response to high intensity blue light, across the 34-h constant routine protocol (n = 12).). Dark bars correspond to the average timing of habitual sleep episodes (biological night). Circadian time 0 corresponds to Dim Light Melatonin Onset (DLMOn, mean ≃ 21:30). **A.** Combined model (sum of linear and sinusoidal components) applied to normalised data (mean ± SD; R² = 0.83). **B.** No linear component (R² = 0.04; p = 0.41). **C.** Sinusoidal component (R² = 0.83). Maximum pupil constriction follows a circadian rhythm with a maximum of sensitivity to high intensity blue light at 10:00.

**Supplementary figure 2:**
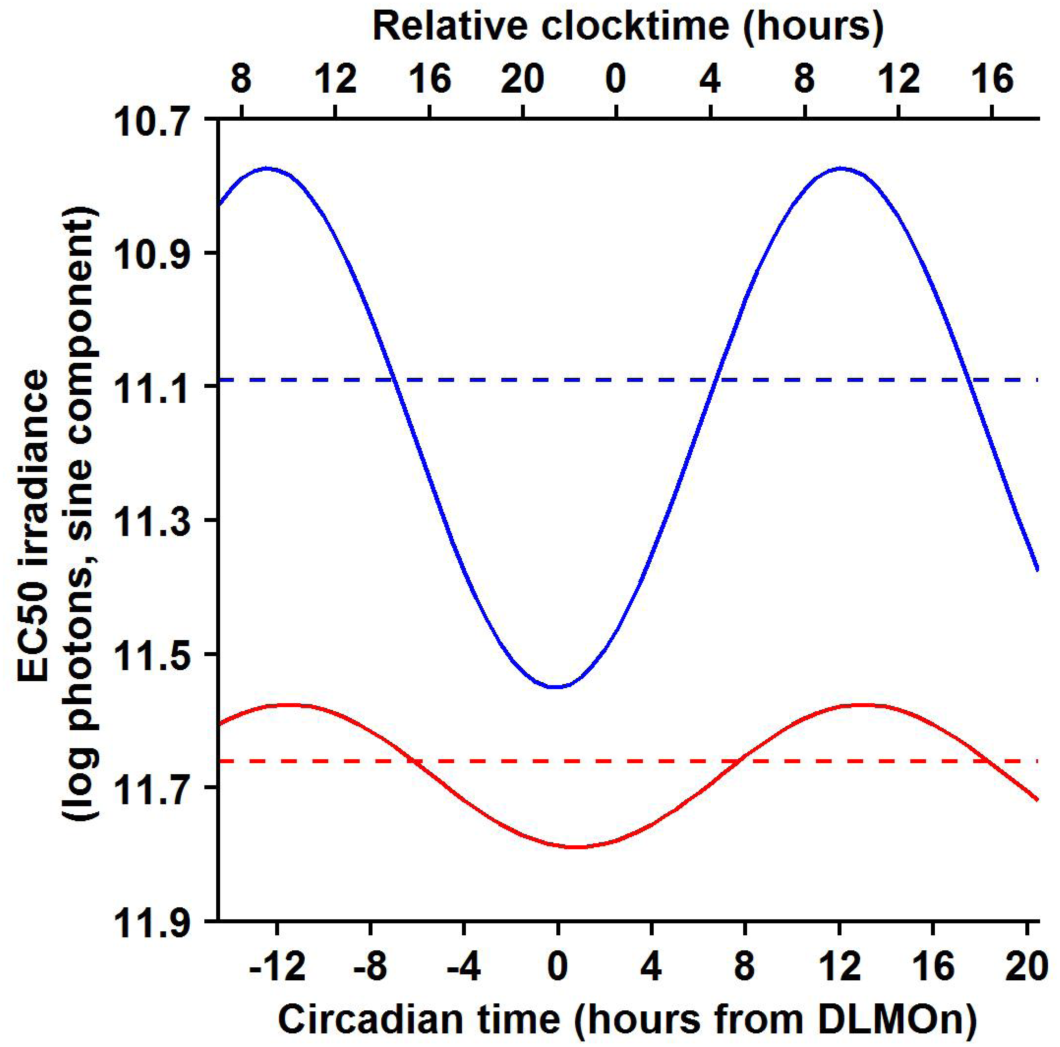
Circadian rhythms of pupillary light reflex in response to low intensity blue, and red light across the 34-h constant routine protocol (n = 12). Circadian time 0 corresponds to Dim Light Melatonin Onset (DLMOn, mean ≃ 21:30). The dotted lines represent the mean sensitivity: 11.66 log photons for red light (red curve) and 11.09 log photons for blue light (blue curve). The maximum pupil response (minimum EC50 value) occurs at 10:00 for red light and at 9:00 for blue light.

**Supplementary figure 3:**
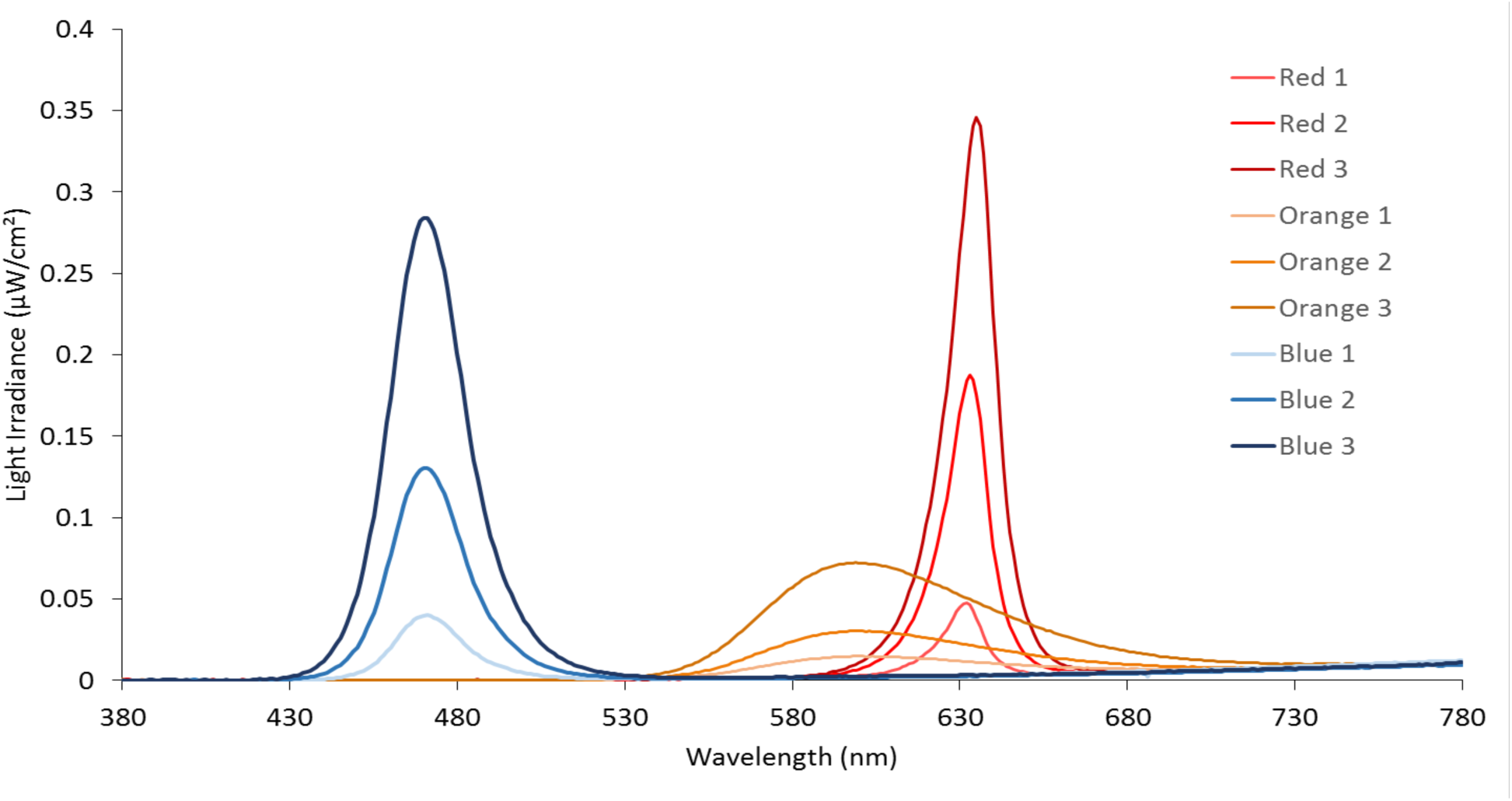
Light spectra for the 9 light stimulations used to induce discomfort glare. The peak of blue light is at 470 nm, orange light at 600 nm and red light at 635 nm. Red 1, Orange 1 and Blue 1 correspond to the lowest irradiance: 12.85 log photons. Red 2, Orange 2 and Blue 2 correspond to the middle irradiance: 13.11 log photons. Red 3, Orange 3 and Blue 3 correspond to the highest irradiance: 13.4 log photons.

**Supplementary figure 4:**
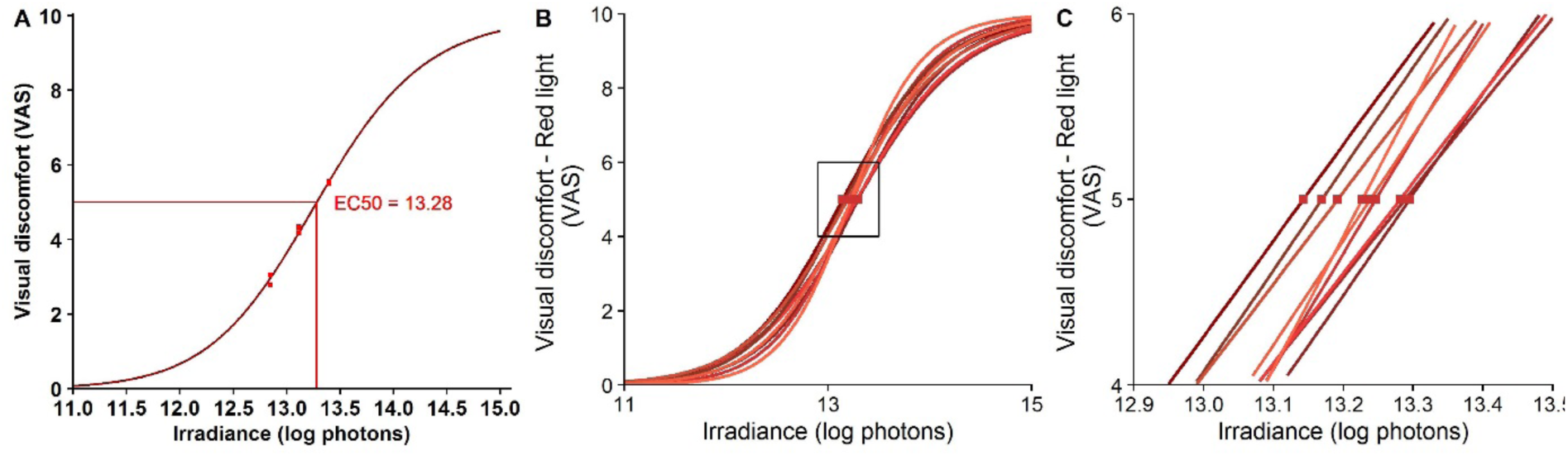
Discomfort glare Intensity Response Curve (IRC) in response to red light. A. Sigmoidal regression at circadian time -1 (∼ 20:30), calculated on a 4-hour time epoch (18:30 – 22:30), corresponding to 2 evaluations of discomfort glare to each of the 3 irradiances (12.85, 13.11 and 13.4 log photons), providing 6 points on the regression curve. The EC_50_ value, corresponding to a light irradiance inducing a discomfort glare of 5/10, is extracted from the sigmoidal regression (here EC_50_ = 13.28 log[irradiance]). **B.** Sigmoidal regression at each of the 4-hour time epochs of the constant routine protocol. The 9 IRCs allow the extraction of 9 EC_50_ values. **C.** Zoom of graph B showing the 9 EC_50_ values that were extracted from these curves and plotted in figure 1A.

**Supplementary figure 5:**
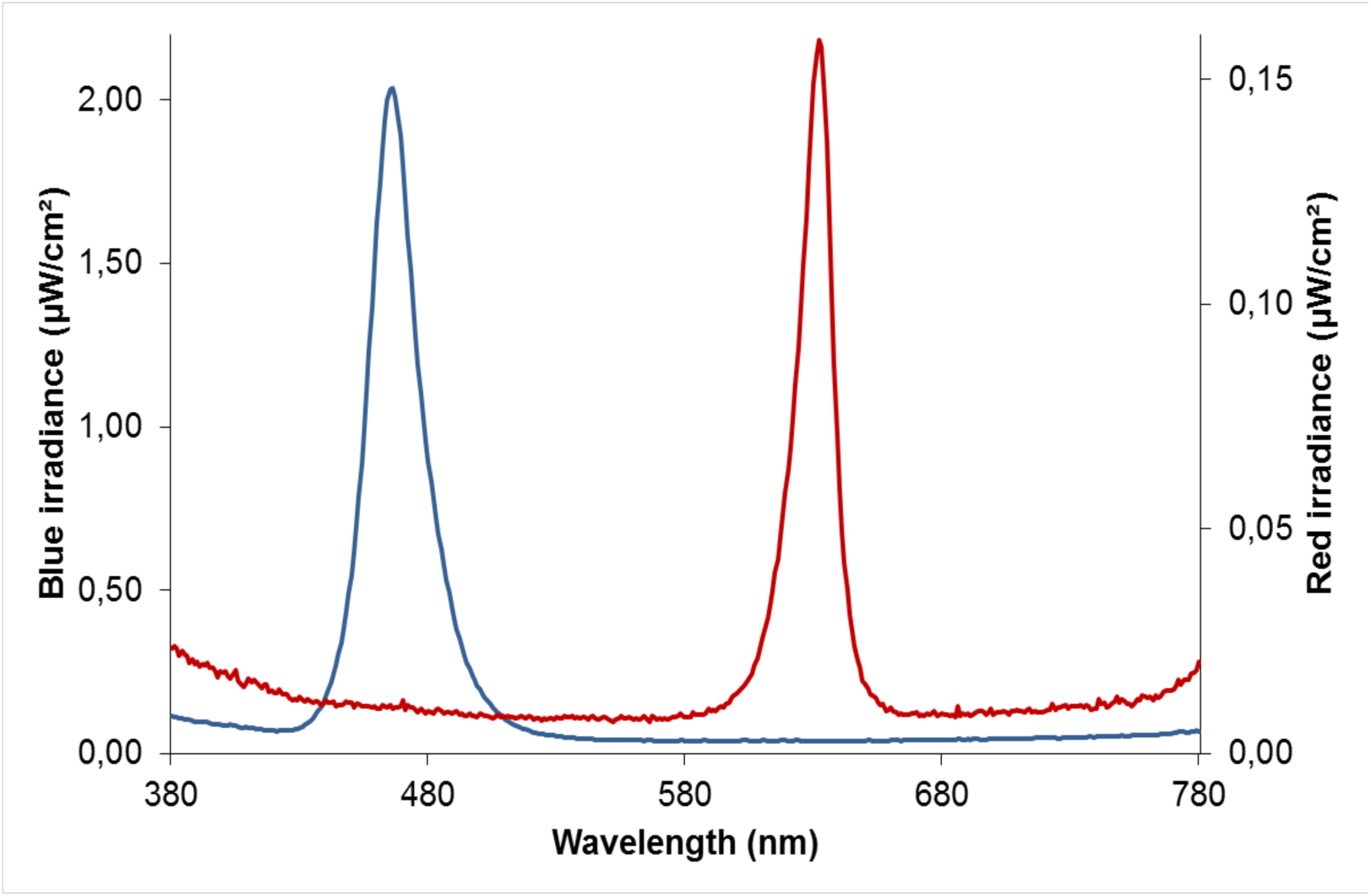
Light spectra for the blue and red light exposures used to induce pupillary light reflex. Peak of blue light is at 465 nm and peak of red light at 630 nm, and the full-width at half of the maximum (FWHM) are respectively 23 and 15 nm. The curves represent the highest irradiances used for blue and red light exposures.

**Supplementary table 1:**
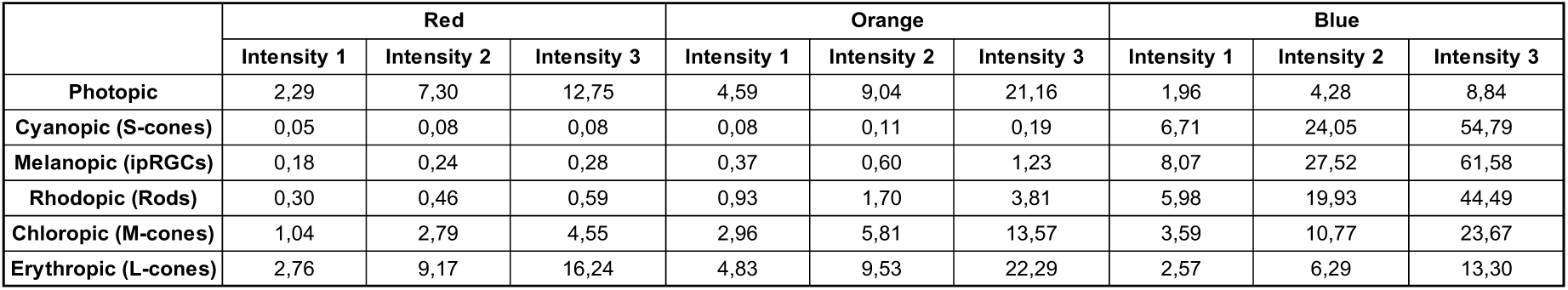
Distribution of photopic and alpha-opic lux content of each of the 9 light exposures.

|  | Red |  |  | Orange |  |  | Blue |  |  |
| --- | --- | --- | --- | --- | --- | --- | --- | --- | --- |
|  | Intensity 1 | Intensity 2 | Intensity 3 | Intensity 1 | Intensity 2 | Intensity 3 | Intensity 1 | Intensity 2 | Intensity 3 |
| Photopic | 2,29 | 7,30 | 12,75 | 4,59 | 9,04 | 21,16 | 1,96 | 4,28 | 8,84 |
| Cyanopic (S-cones) | 0,05 | 0,08 | 0,08 | 0,08 | 0,11 | 0,19 | 6,71 | 24,05 | 54,79 |
| Melanopic (ipRGCs) | 0,18 | 0,24 | 0,28 | 0,37 | 0,60 | 1,23 | 8,07 | 27,52 | 61,58 |
| Rhodopic (Rods) | 0,30 | 0,46 | 0,59 | 0,93 | 1,70 | 3,81 | 5,98 | 19,93 | 44,49 |
| Chloropic (M-cones) | 1,04 | 2,79 | 4,55 | 2,96 | 5,81 | 13,57 | 3,59 | 10,77 | 23,67 |
| Erythropic (L-cones) | 2,76 | 9,17 | 16,24 | 4,83 | 9,53 | 22,29 | 2,57 | 6,29 | 13,30 |

